# Environmental variables and species traits as drivers of wild bee pollination in intensive agroecosystems -a metabarcoding approach

**DOI:** 10.1101/2023.03.20.533466

**Authors:** Marina Querejeta, Lorène Marchal, Paul Pfeiffer, Marilyn Roncoroni, Vincent Bretagnolle, Sabrina Gaba, Stéphane Boyer

**Affiliations:** Institut de Recherche sur la Biologie de l’Insecte, UMR 7261, CNRS-Université de Tours, Tours, France; Department of Functional Biology, University of Oviedo, Asturias, Spain; UMR UREP, INRAE, 5 Chemin de Beaulieu, 63000 Clermont-Ferrand, France; CEBC, UMR7372, CNRS and La Rochelle Université, 405 Route de Prisse la Charrière, 79360 Villiers-en-Bois, France; LTSER « Zone Atelier Plaine & Val de Sèvre », F-79360 Villiers-en-Bois, France; USC 1339 Résilience, Centre d’Études Biologiques de Chizé, INRAE, F-79360 Villiers-en-Bois, France

**Keywords:** Wild bees, sunflower, pollen DNA metabarcoding, pollination, biodiversity

## Abstract

Wild bees are known to be efficient pollinators of wild plants and cultivated crops and they are essential ecosystem service providers. However, wild bee populations have been suffering from significant declines in the last decades mainly due to the use of agrochemicals. Within this framework, we aimed to characterize wild bees pollination spectrum (i.e. the community of pollinated flowering plants) in intensive agroecosystems, and describe the environmental variables and wild bee species traits influencing the pollination. To do this, we conducted metabarcoding analyses of pollen loads from wild bees collected in sunflower crops in the French region of Nouvelle-Aquitaine. Our study revealed that wild bees visited flowering plants corresponding to 231 different Operational Taxonomic Units, classified in 38 families of which Asteraceae, Brassicaceae and Apiaceae were the most visited and more than 90% of the visited taxa turned out to be wild flowers. We also analysed the potential effect of environmental variables and wild bee species traits in governing their choice of pollinated plants. The community composition of pollinated plants varied depending on the flowering stages of the sunflower and the farming system. Our results also show that pollination niche breadth (alpha diversity) varied depending on the flowering stages of the sunflower but was not different between organic and conventional farming systems.

Regarding wild bee species traits, the community composition of pollinated plants varied in relation to wild bees body sizes and, sociality levels. Our results are consistent with previous studies, suggesting that solitary bees are more specialists when it comes to flower selection than social bees, which are more generalist. The metabarcoding of pollen loads enabled us to draw a global picture of plant-wild bee interactions in an intensive agroecosystem. Our findings support the hypothesis that a higher diversity of weeds may increase wild bee diversity in intensive agroecosystems.

## 1 Introduction

Wild bees (monophyletic group Apiformes, clade Antophila, non-*Apis* bees) are a diverse group from a species-richness, phylogenetical and behavioral perspectives, and comprises around 18,000 species worldwide divided in seven different families (Michener, 2000; Winfree, 2010). This diversity has enabled wild bees to colonize a wide variety of ecosystems, being present in every continent of the planet, with the exception of Antarctica (Danforth, 2007). Communities of wild bees are among the most efficient pollinators (Garibaldi *et al*., 2013) and they are responsible for 20% of the global pollination services (Losey and Vaughan, 2006; Sánchez-Bayo and Wyckhuys, 2019). They are, thus, providers of an essential ecosystem service given that at least, one third of our food, including animal origin food, depends on animal-pollinated (usually bee-pollinated) crops (McGregor, 1976; Klein *et al*., 2007; Aizen *et al*., 2009). Be that as it may, several species are food specialists with narrow foraging ranges and nesting habitat resources (Roulston and Goodell, 2011; Sánchez-Bayo and Wyckhuys, 2019), which makes them vulnerable to environmental changes. Indeed, in the course of the last century wild bee populations have suffered widespread declines (Zattara and Aizen, 2021) alongside the rest of insect taxa (Basset *et al*., 2012; Leather, 2017; Basset and Lamarre, 2019; Cardoso *et al*., 2020; Didham *et al*., 2020). One of the main causes of the decline of wild bee populations is the loss of suitable nesting sites and foraging habitats (Foley *et al*., 2005; Steffan-Dewenter, Potts and Packer, 2005; Brown and Paxton, 2009), which has been reported to be linked to intensive agriculture, and the use of agrochemicals within agricultural landscapes (Basset *et al*., 2012; Leather, 2017; Basset and Lamarre, 2019; Brühl and Zaller, 2019; Cardoso *et al*., 2020; Didham *et al*., 2020).

Agrochemicals have direct toxic effects on insects but, also indirect effects such as habitat alteration (Hayes *et al*., 2017), which may modify the abundance and diversity of weeds (wildflowers) within and around crops, thereby altering the availability of floral resources for pollinators (Potts *et al*., 2010). These threats can cause the disappearance of rare pollinator species and lead to the homogenization of natural plant and pollinator communities (Hallmann *et al*., 2017; Seibold *et al*., 2019), which in turn affects the functioning of agroecosystems, rendering them more vulnerable to perturbations such as droughts or pest invasions (Silverman and Brightwell, 2008; Roy *et al*., 2016). There is therefore an urge to find solutions to halt the extinctions that insect populations are currently facing (Basset and Lamarre, 2019; Sánchez-Bayo and Wyckhuys, 2019). In the scenario of extinction of pollinators, estimations of high economical losses have been made (Gallai *et al*., 2009; Sandhu *et al*., 2016). Also, potential failures in pollination would decrease long-term survival of plants (Biesmeijer *et al*., 2006).

Conventionally, pollination is studied through observational approaches (Arizaga *et al*., 2000; dos Santos and Wittmann, 2000; [1] [2], which require significant efforts to obtain only limited information [3], as the focus is mainly on the plant being pollinated. In the last decade, the boom of high throughput DNA sequencing technologies and, especially, the use of environmental DNA (eDNA) metabarcoding has enabled scientists to acquire more information about community composition in an efficient and cost-effective manner for large scale surveys [4].eDNA has become a valuable tool to monitor biodiversity and interactions [5] [6] [7] in a wide variety of ecosystems, including plant-pollinator interactions within agroecosystems [8] [9] [10] [11]. One of the strengths of eDNA is the fact that it provides information about composition of ecological communities. Exploring pollination and acquiring knowledge regarding the quantity of pollen carried by pollinators is key to provide an accurate view of the global pollination picture than those estimations obtained through observational approaches [12] [13] [14].In fa ct, molecular-based identification techniques of pollen have shown to identify a higher number of plant taxa with greater taxonomic resolution than observational approaches and it does not rely on taxonomy experts [15] [16]. Within this framework and with the final objective of integrating the knowledge acquired into conservation programs, we developed an ITS metabarcoding approach on pollen loads to describe the pollination spectrum of wild bees in sunflower crops, as a model for intensive agroecosystems. We also aim to describe which environmental variables govern their choices of pollinated plants. Lastly, we tested the impact of two wild bee species traits, body size and social level (social vs. solitary), not only on pollinated flowering plant community composition but also on pollination niche breadth (diversity of pollinated plants). Acquiring knowledge of flowering plants pollinated by wild bees’ communities (hereafter called pollination spectrum) and the drivers of this pollination process might help understanding pollinator-plant interactions in agroecosystems and potentially inform effective ecological intensification approaches to improve the conservation status of wild bees.

## 2 Material and Methods

### 2.1 Wild bee collection and characterization

We captured wild bees (non-*Apis* species) in the LTSER Zone Atelier Plaine & Val de Sèvre (SW France, Nouvelle-Aquitaine Region) (Bretagnolle *et al*., 2018), in 25 sunflower fields with a maximum of 10 individuals per field. A total of 212 specimens were collected, exclusively during the pre-flowering (7^th^ – 9^th^ July 2020) and flowering (17^th^ – 22^nd^ July 2020) periods (Table S1A; S1B). Wild bees were collected on any flowering plant within the field and its edges using either a sweep net or directly with a 25 ml sampling tube. The specimens were then placed in a cooled ice box for transportation to the laboratory, where they were stored at -20^◦^C until subsequent processing.

Together with wild bee specimen collection, we took records of the following environmental variables in each crop: percentage of ground covered by weeds in the field, sunflower crop flowering period (pre-flowering or flowering), height of the sunflower (cm), percentage of sunflower flowering and farming system (organic or conventional). Percentage of weeds and height of sunflower were categorized in 4 quartiles.

Wild bee specimens were identified morphologically to species level. Wild bees were classified by two species traits: size (categorized as T1 to T4) and sociality level (social or solitary). Moreover, we also calculated abundance and species richness of wild bees for each environmental variable explained above: sunflower flowering period, farming system, percentage of weeds and height of sunflower. Finally, we measured the differences in wild bee community composition (Whittaker index) or the beta diversity between the levels of the four environmental variables, which ranged from 0 (no species turn over) to 1 (every sample holds a unique set of species), using the *vegan* R package [17].

### 2.2 Wild bee sample processing, pollen DNA library preparation and flowering plant sequencing

Prior to DNA extraction, all wild bee specimens were individually washed to recover the pollen from their cuticle and wings following a method developed by [18]. Following DNA extraction, an ITS2 metabarcoding library was prepared. To that purpose, we used the primer pair (5-ATGCGATACTTGGTGTGAAT-3) and ITS4R (5-TCCTCCGCTTATTGATATGC-3) (White et al., 1990; [19] modified for high-throughput sequencing (HTS), we attached Nextera XT adaptors through a second amplification and we performed a final equimolar pooling (see detailed protocol in Supplementary information). This library was sequenced in Illumina MiSeq using V2 chemistry (250 × 300 bp, 500 cycles) in the Sequencing Center within the Biozentrum of the Ludwig-Maximilian-University in Munich (Germany).

### 2.3 ITS metabarcoding library filtering and taxonomic assignment

The raw library was filtered using a standard toolbox of software. The expected sequencing depth was around 78,000 reads per sample. The quality of the library was checked using *FastQC* (https://www.bioinformatics.babraham.ac.uk/projects/fastqc/), primers were trimmed using *cutadapt* [20] and reads were merged with *PEAR* [21]. The remaining filtering and Operational Taxonomic Unit (OTU) clustering (97% of cut-off threshold) was performed using *vsearch v2*.*8*.*2* [22]. At this point, we had produced an OTU table. When possible, OTUs were classified until species level using *rjson* [23], taxize R packages (Chamberlain and Szöcs, 2013) and a phylogenetic confirmation through IQ-TREE [24] (see detailed bioinformatic protocol in Supplementary information). All samples with less than 1,000 reads were discarded for subsequent analyses. Flowering plants species (OTUs) for which the DNA was detected in our samples, were considered to have been visited by the bees and therefore potentially pollinated.

### 2.4 Flowering plant biodiversity data analysis

At this point, only the wild bee samples with a minimum of 1000 reads were kept for subsequent analyses. Thus, we estimated the completeness of the sampling by computing a rarefaction curve and individual rarefaction curves for each of the environmental variable and species trait studied using R package *vegan* (Oksanen *et al*., 2013). As another measure of quality, we calculated the number of OTUs detected in each sample and averaged this number at the population level to determine the number of pollinated plants detected per wild bee specimen. We used the multivariate BIO-ENV procedure (Clarke and Ainsworth, 1993), which is a dissimilarity-based method to identify the set of environmental variables that are most correlated to the OTU table, and thus best explain the bee pollination spectrum. The four environmental variables for which we had records were included in the BIO-ENV procedure: flowering period, farming system, percentage of weeds and height of sunflower.

The flowering plants pollinated by wild bees on sunflower crops were described using two different biodiversity metrics, Relative Read Abundance (RRA) and Frequency of Occurrence (FOO). RRA refers to the proportion of reads obtained per OTU in each sample, each variable or for the full dataset and FOO informs about the number of samples (counts) in which an OTU is present (Deagle *et al*., 2019). The calculation of both metrics, RRA and FOO, for the whole sampling and for each the selected variables’ levels was calculated using customized scripts based on R package *dplyr* (Wickham *et al*., 2021).

The importance of environmental variables and wild bee species trait in the pollination spectrum was inferred by fitting them onto an unconstrained ordination and applying *envfit* analysis [17]. Alpha diversity (Exponential of Shannon Index, q = 1) of pollinated plants was calculated using *hilldiv* R package [25].

To test the statistical significance of the differences in alpha diversity, we computed a Kruskal-Wallis (K-W) rank sum test and the post-hoc Dunn test in the case of variables with more than two levels, using *stats* [26]and *dunn*.*test* [27])R packages, respectively. Beta diversity (community composition) was calculated using an unconstrained ordination through a principal coordinate analysis (PCoA) and a constrained ordination through canonical analysis of principal coordinates (CAP). Both these analyses were based on Bray-Curtis distance matrix to account for differences in abundance, applying the *phyloseq* R package [28]. Statistical significance of both ordinations was tested through a non-parametric multivariate analysis or PERMANOVA (999 iterations) and constrained CAP permutation test, respectively [17]. For the case of these two permutational analyses, aside from the main effects, the interaction between the environmental variables and the species traits was also tested.

We explored the effects of environmental variables and species traits selected, together with the interaction between them through the computation of a negative binomial generalized linear model (GLM), using a nested model and applying a log link function (999 iterations) using *mvabund* R package [29].

### 2.5 Differentially abundant flowering plant

For each of the selected environmental variables and species traits, we identified the differentially abundant flowering plant OTUs by applying a *DESeq* negative binomial Wald test using *DESeq2* R package [30], that is to say, the OTUs, which show a significantly higher abundance in a particular factor within the environmental variables or species traits. *DESeq2* analysis is a procedure originally designed for differential expression analyses of genes, however, it is widely used in count/abundance data. The identified plant species (OTUs) were considered as indicator species of each factor of the environmental variable or species trait and were further analyzed. Significance of differential abundance among levels of variables was tested through Kruskal-Wallis test, and the RRA of significantly abundant species visualized through a hierarchically clustered (complete linkage method) using the *pheatmap* R package [31].

### 2.6 Plant-wild bee ecological networks

For each species trait (body size and sociality level), we constructed bipartite networks, based on the RRA of flowering plant OTUs using the *bipartite* R package [32]. OTUs representing less than 1% of the reads were discarded from the network to obtain a better visualization. All the biodiversity analyses, *DESeq2* analysis and bipartite networks were done using R version 3.6.1 [33].

## 3. Results

### 3.1 Characterization of wild bee communities

We collected 212 wild bees from sunflower crops, which were identified at species level and corresponded to 33 different species (Table S2; Fig. S1). The most abundant wild bee species was the social sharp-collared furrow bee (*Lassioglossum malachurum*), while the yellow-legged mining bee (*Andrena flavipes*) was the most abundant among the solitary bees (Fig. S1A). The wild bees included in this study belonged to four families (out of seven described): Andrenidae, Colletidae, Halictidae and Megachilidae (Fig. S1A), with the majority belonging to Halictidae. The overall distribution of wild bee families was similar among both flowering periods and farming systems (Figs. S1B and S1C).

Wild bee species richness did not show important differences among the selected environmental variables, flowering period and farming system. Between the two flowering periods the Whittaker index (beta diversity) was 0.387 and between the two farming systems 0.36 (values ranged from 0 to 1) (Figs. S1B and S1C).

### 3.2 Description of library quality and wild bee’s pollination spectrum

Out of the 212 pollen samples, 185 passed the sequencing depth filter (> 1,000 reads) and produced a total of 15,153,994 reads (i.e., an average 71,484.1 reads per sample). Reads assigned to sunflower *Helianthus annus* (10,707 reads, 1.18%) were found in the extraction control and subtracted from all the samples. According to the rarefaction curves (Fig. S2A), the sequencing quality was adequate to study the plants pollinated by wild bees in sunflower crops. All rarefaction curves reached a plateau (Fig. S2B) and we detected an estimated 90.53% of the OTUs of plant species pollinated by wild bees. The mean number of plant OTUs detected per wild bee sample was 12.12. A total of 269 OTUs were detected, of which 231 were kept for subsequent analyses as they corresponded to flowering plants (class Magnoliopsida) and 150 were identified at species level (Table S2). Out of the 38 OTUs discarded, one was assigned to a fungus (class Ascomycota), 13 to uncultured eukaryotes and the 24 OTUs remaining did not reach our quality thresholds (percentage of identity > 75% and query coverage > 60%).

The flowering plant community was taxonomically classified in 21 orders, 38 families, 108 genera and 153 different species within the 231 OTUs (Figs 1A and 1B). Out of the 38 families detected, 8 represented more than 1% of the filtered reads and 29 were present in more than 1% of the samples (Fig. 1A). Asteraceae (RRA = 30.85%) and Brassicaceae (RRA = 28.64%) were the most abundant families followed by Apiaceae (RRA = 13.78%) and Convolvulaceae (RRA = 8.68%) (Fig. 1A). With regards to prevalence, the most common families (highest FOO) were Asteraceae (FOO = 72.29%), Brassicaceae (FOO = 57.58%), Apiaceae (FOO = 53.25%), Plantaginaceae (FOO = 34.63%) and Rosaceae (FOO = 29%) (Table S2). At species level (Fig. 1B, Table S2), white mustard *Sinapis arvensis* (RRA = 21.47%) was the most abundant, followed by wild carrot *Daucus carota* (RRA = 12.85%), hawkweed oxtongue *Picris hieracioides* (RRA = 8.84%) and field bindweed *Convolvulus arvensis* (RRA = 8.36%) (Fig. S3). In terms of occurrence in samples (Table S2), *D. carota* (FOO = 48.92%) and *S. arvensis* (FOO = 44.16%) were the most common species followed by the stinking chamomile *Anthemis cotula* (FOO = 37.66%) and the common sowthistle *Sonchus oleraceaus* (FOO = 28.57%). Globally, 184 OTUs (RRA = 93.1%) corresponded to wild plants while 47 OTUs (RRA = 6.9%) were crop or ornamental plants (Fig. 1C).

**Figure 1:**
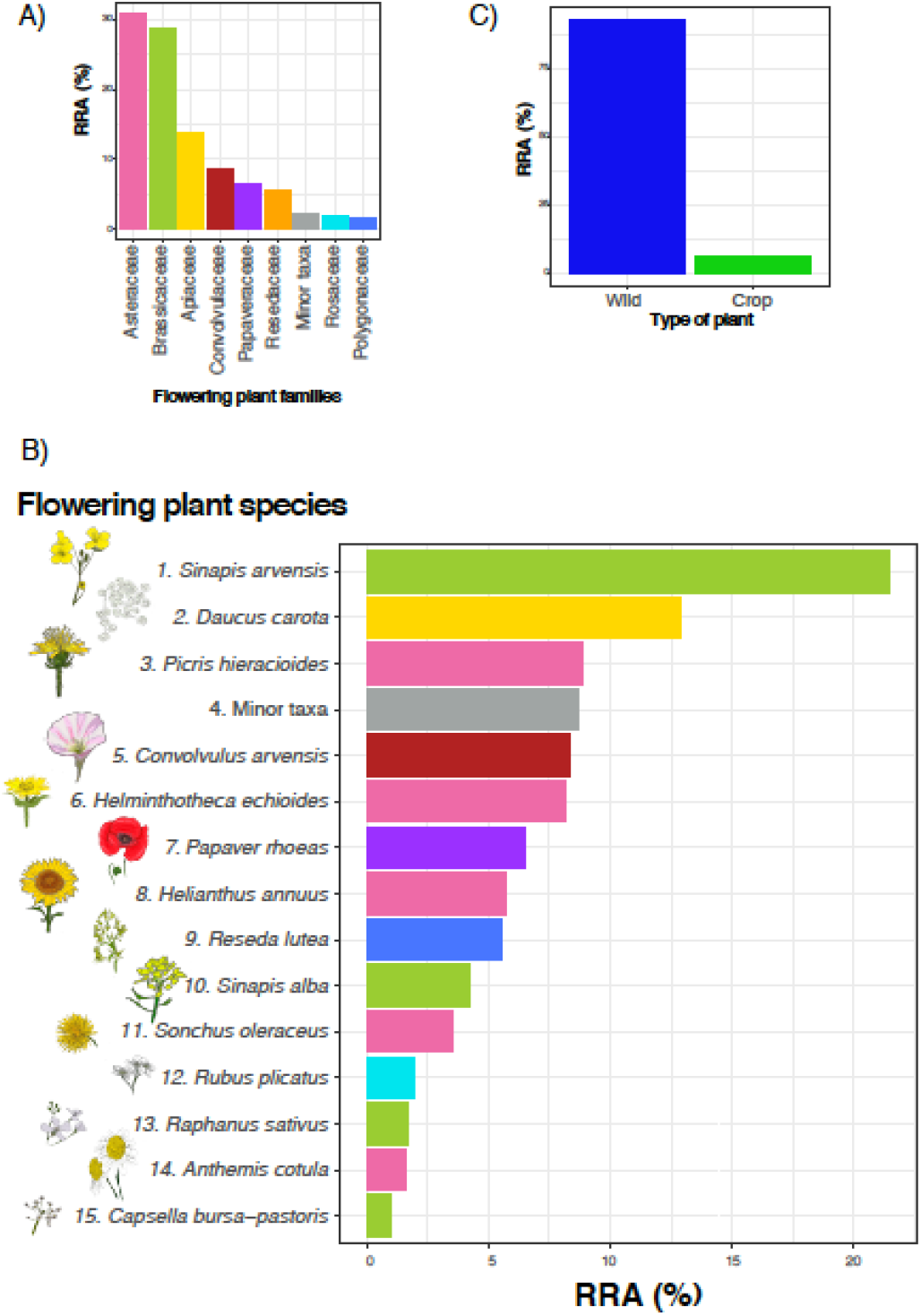
Relative Read Abundance (RRA) of the flowering plant community pollinated by wild bees in sunflower crops in the study area of Plaine et Val de Sèvre. A) assigned until family level (families comprising less than 1% of reads were grouped in the category “Minor taxa”), B) assigned until species level (species comprising less than 1% of reads were grouped in the category “Minor taxa”) and colored by family, C) categorized by type of plant (wild or crop plant).

### 3.3 Environmental variables and species traits driving change in pollination spectrum

According to the BIO-ENV procedure, the flowering period (pre-flowering vs flowering) and the farming system (organic vs conventional) (r = 0.14) were the environmental variables best correlated to the OTU table. Deeper investigation through *envfit* showed that these two environmental variables as well as the two species traits, body size and sociality level, influenced the pollination spectrum the most (prand < 0.001). According to *envfit* outcome, the most important variable explaining the variation in the pollination spectrum was the size of wild bees (R^2^ = 0.218), followed by farming system (R^2^ = 0.1469), social behavior (R^2^ = 0.0941) and flowering period (R^2^ = 0.0744).

In terms of pollinated plant community composition (based on RRA), there were a number of differences between the two flowering periods and the two farming systems (Fig. 2A and 2B; detailed analysis in Supplementary information).

**Figure 2.**
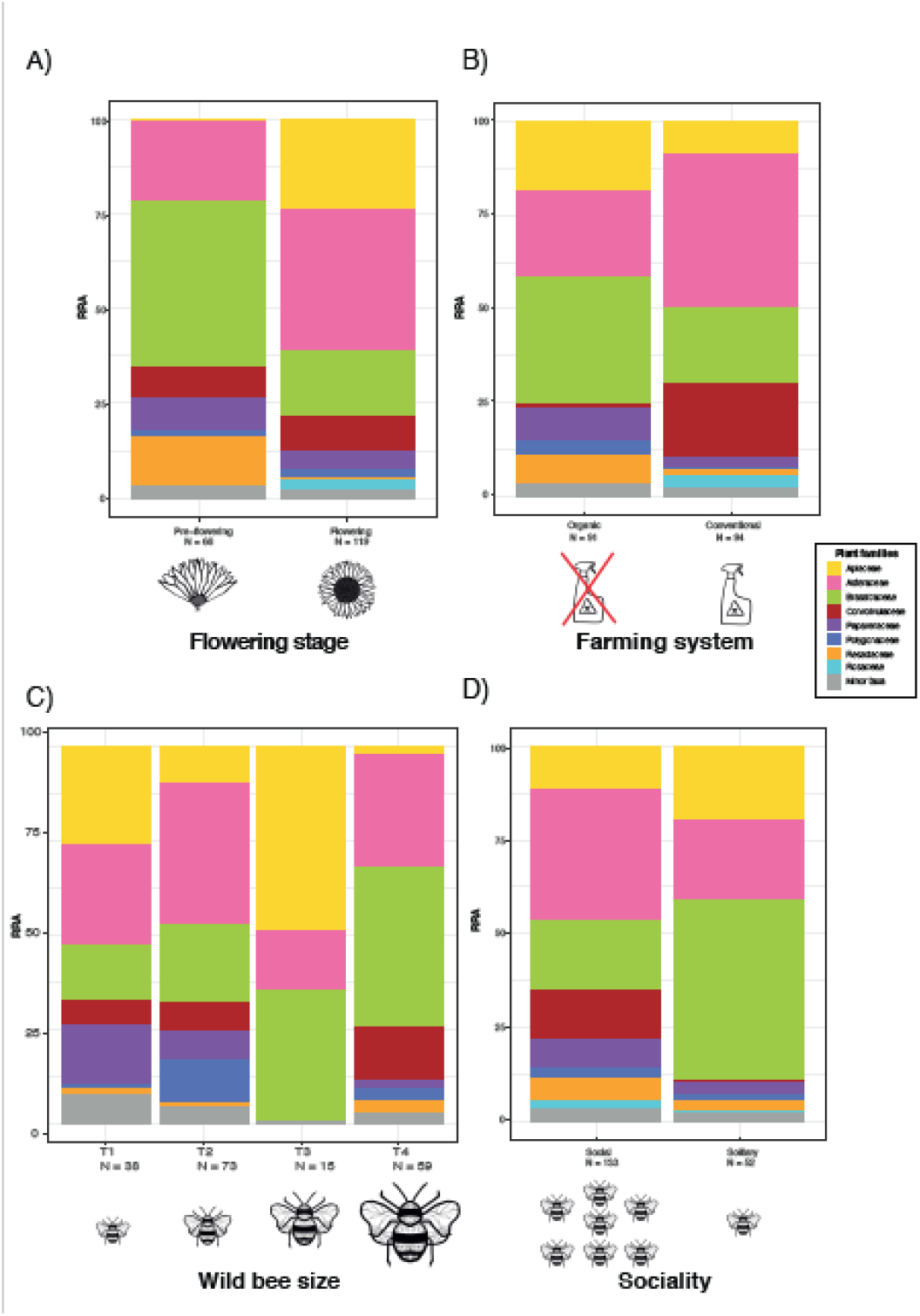
Stacked bar plots of the taxonomic composition of the flowering plants pollinated by wild bees (RRA) at family level among in relation to A) stages of the crop (pre-flowering vs flowering), B) type of agriculture (organic vs intensive), C) Wild bee size categories (T1-T4) and D) sociality (social vs solitary bees).

With regards to environmental variables, we found a significant difference in alpha diversity between both flowering periods with higher diversity of plants pollinated by bees before flowering of the crop compared to the flowering period (K-W Chi^2^ = 4.7447, p-value = 0.02939) (Table S4; Fig. S4A). In contrast, we did not find significant differences in pollinated plant alpha diversity when comparing the two farming systems (K-W Chi^2^= 0.3358, p-value = 0.5623) (Table S4; Fig. S4B).

Beta diversity between the two different flowering periods and the farming systems showed a separation between the communities of pollinated plant but also a certain degree of overlap in both cases on the PCoA (Figs. S4A and S4B). PERMANOVAs indicated that both the flowering period and the farming system had a significant effect in the pollination spectrum of wild bees (R^2^_period_ = 0.03159, prand < 0.001; R^2^_farming_ = 0.03234, prand < 0.001) (Table S4).

Deeper constrained investigation of beta diversity among both levels of each environmental variables was visualized through CAP (Figs. S6A and S6B). Permutation test for CAP confirmed that the differences visualized were significant (Chi^2^_period_ = 0.379, prand < 0.001; Chi^2^_farming_=0.3562, prand < 0.001) (Table S4).

As for species traits (body size and sociality level), differences in alpha diversity between the four wild bee size categories were not significant according to Kruskal-Wallis test (K-W Chi^2^ = 6.58, p-value = 0.09). Regarding sociality level, we found significant differences in alpha diversity between social levels (K-W Chi^2^= 3.9292, p-value = 0.04745) (Table S4; Fig. S4D and Fig. S7) and social wild bees showed a more diverse pollination spectrum than the solitary community.

Regarding community composition (beta diversity) of the species traits, body size and sociality level, was visualized through the unconstrained PCoA, showing a certain degree of dissimilarity among communities (Figs. S5C and S5D). PERMANOVAs showed a significant effect of, both, the four wild bee body size categories and the two sociality levels on the pollination spectrum of wild bees (R^2size^ = 0.04865, prand < 0.001; R^2^_social_ = 0.01764, prand < 0.001) (Table S4).

Further constrained investigation of beta diversity was visualized through CAP (Figs. 4C and 4D). Permutation test for CAP permutation test confirmed the statistical significance of the visualized differences (Chi^2^_size_ = 0.7709, prand < 0.001; Chi^2^_social_ = 0.2243, prand < 0.001) (Table S4). In the case of the wild bee size, CAP shows an overlap in community composition with the biggest separation between T1 and T3 (Fig. S6D). In the case of bee sociality level (Fig. S6C), we detected a clear separation between social and solitary bees but, also, a certain degree of overlap between both sociality levels. The interaction between the two environmental variables and the two species traits was significant in the case of PERMANOVA (R^2^_interaction_ = 0.2717, prand < 0.001) and CAP permutational analysis Chi^2^_interaction_ = 4.6465, prand < 0.001) (Table S4).

Effects of each environmental variable and species trait (main effects) and interaction on the pollination spectrum of wild bees showed up to be significant according to GLM (Dev_period_ = 797.1, prand < 0.001; Dev_farming_ = 746.3, prand < 0.001; Dev_size_ = 1235, prand < 0.001; Dev_size_ = 436.8, prand < 0.002; Dev_interaction_ = 4924, prand < 0.00) (Table S4).

### 3.4 Flowering plant species as indicators of environmental conditions and wild bee traits

*DESeq2* and K-W detected 15 plant species (25 OTUs), which were differentially abundant in relation to environmental conditions or wild bee species traits (Table S3). Among these 15 plant species, those showing the highest RRA for a specific environmental condition or wild bee species trait are considered as indicator species. *Reseda lutea* was an indicator species during pre-flowering while *D. carota* was an indicator during flowering of sunflower (Fig. 3A). Concerning the farming system (Fig. 3B), *D. carota* was also an indicator of organic crops while *C. arvensis* turned out to be an indicator of conventional crops. Among the wild bee size categories (Fig. 3C), the common poppy *Papaver rhoeas* was preferred by small bees (T1 category), while T2 bees preferred *Hypochaeris sp*. and T3 *P. hieracioides*. No plant indicator species was detected for the large wild bee category T4. As for social levels (Fig. 3D), social bees preferred *C. arvensis* while *Convolvulus sp*. was preferred by solitary bees.

**Figure 2:**
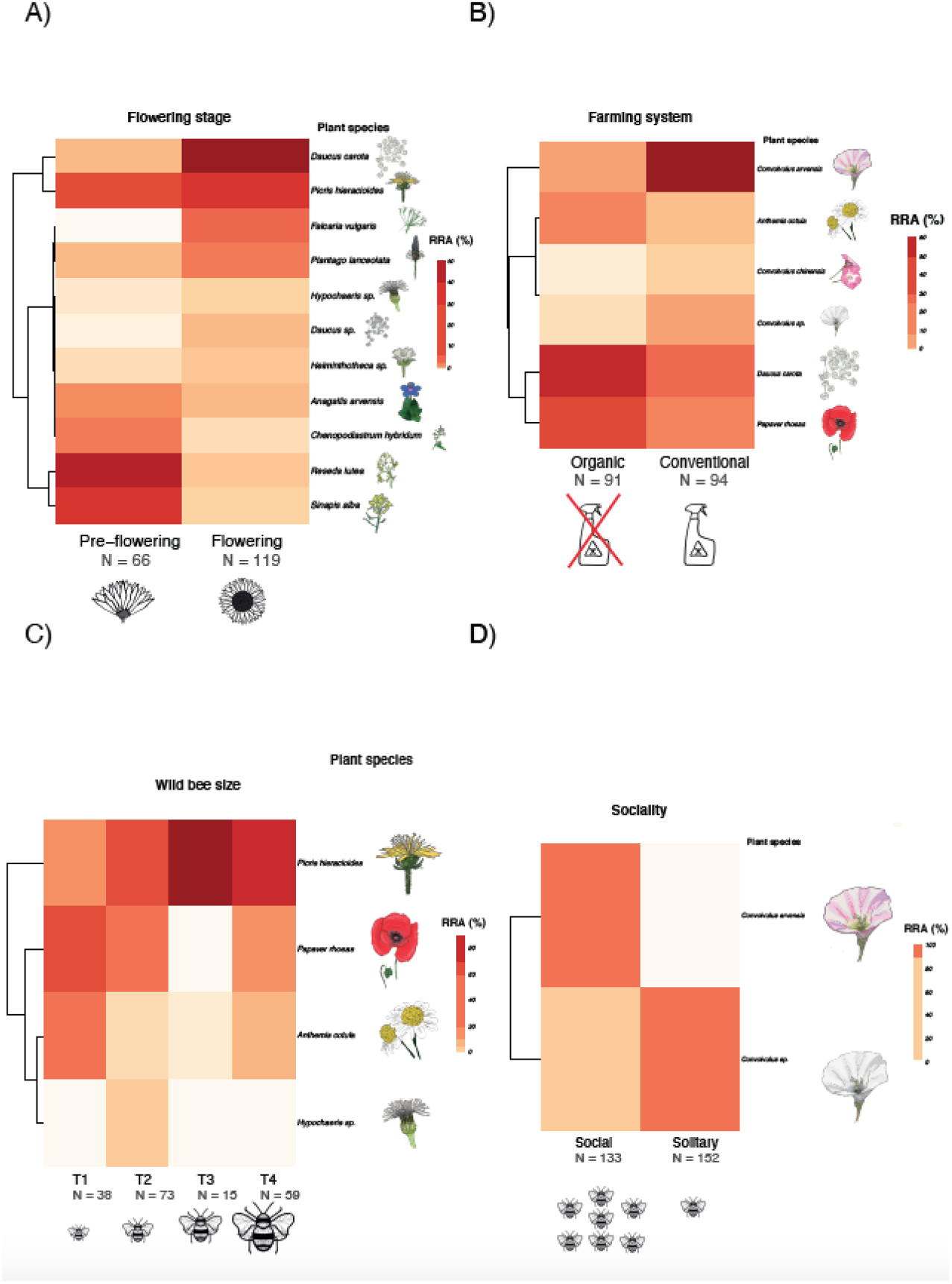
Clustered heatmap of RRA from differentially abundant OTUs clustered by species according to DESeq2 analysis and Kruskal-Wallis test in relation to A) stages of the crop (pre-flowering vs flowering), B) type of agriculture (organic vs intensive), C) Wild bee size categories (T1-T4) and D) sociality (social vs solitary bees).

### 3.5 Plant-pollinator ecological networks

Bipartite ecological networks showed wide overlap in the pollination spectrum of solitary and social bees (Fig. 4A). Five plant species, however, were exclusively pollinated by social bees (*Cirsium arvense, Crepis capillaris, Polygonum boreale, C. arvensis, A. cotula*), while only one plant species (*Capsella bursa-pastoris*) was exclusively pollinated by solitary bees. With regards to bee body size (Fig. 4B), eight plant species were exclusively pollinated by one bee size category (*A. cotula, Lactuca sativa, C. capillaris* and *P. boreale* pollinated only by T1 bees; *P. lanceolata, F. vulgaris, C. arvense* were pollinated only by T2 bees; and *C. bursa-pastoris* pollinated only by T3 bees). In contrast, six species of plants provided resources to all four different size categories of bees, these plants included the main crop plant (*H. annuus*) as well as common weeds (*P. hieraciodes, S. oleracerus, D. carota, S. alba, S. arvensis*).

## 4 Discussion

We analysed plant species composition from pollen collected by wild bees that were captured in sunflower crops. Aside from characterizing the pollination spectrum of wild bee communities, this study revealed that wild plants are key for the survival of wild bees even within an intensive agroecosystem and even during sunflower blooming period, as they show an overall preference for wild floral resources. Moreover, despite the overlap in floral resources used by different species of wild bees, we found clear differences in the choice or availability of plant taxa in relation to several environmental variables, such as flowering period and farming system. In addition, the size of wild bees and their sociality appears to have an influence on their pollination spectrum in sunflower crops, together with the interaction of the mixed effect between environmental variables and species traits. However, because of working at community level, there are different sampling sized between the factors of environmental variables and species traits. Therefore, these results should be taken with caution as these sampling bias inherent to the type of study could have an impact, especially in the case of alpha diversity.

### 4.1 Pollen DNA metabarcoding as an efficient tool to describe pollinated plant communities

ITS2 DNA metabarcoding has been used with success to understand trophic interactions in agricultural landscapes [34] [18]. Here, pollen DNA metabarcoding has allowed us to identify plant species visited, and therefore potentially pollinated, by wild bees. This method has enabled us to not only directly detect and identify to species level a high percentage (69%) of the pollination spectrum of the collected wild bees, but also relate this pollination spectrum to environmental variables and wild bee species traits. Such high precision and accuracy would be challenging to reach using pollination observational studies (Arizaga *et al*., 2000; dos Santos and Wittmann, 2000; [1] [2], as these studies are plant-based and produce information only from one plant at each observation. In contrast, our metabarcoding analysis led to the identification of more than 10 plant species per wild bee, which means that, as suggested by previous studies [15] [16], we are discovering additional hidden information compared to conventional methods. On the other hand, metabarcoding approaches are not free of biases. Despite their current biases, the confirmation that our pollen load DNA metabarcoding approach is able to recover such a high-quantity and high-quality information about plant-pollination interactions opens a promising avenue of research as this method requires less effort and less botanical expertise than traditional observation methodologies.

### 4.2 Weeds as essential floral resources for wild bees

Our study illustrated that, even if flowering crops within intensive agricultural landscapes can provide floral resources to pollinators [35], these are usually transient plant taxa, which contribute with an increase of pollination rates only during reduced periods of time [36]. Thus, crop plants do not always overlap with pollination season and many wild pollinators not only need enough number of flowering plants to pollinate but also a continuity in the availability of floral resources across a prolonged period [37]. For instance, sunflower blooming stage lasts around four weeks per year [38],which is a short period to fulfil the requirements of wild bee communities. In contrast, the presence of weeds inside or on the edges of the crops, which are more diverse in terms of flowering periods, sizes and shapes, increases pollination rates of wild bees [39] [40] [41] [42]. This goes in line with our study showing that flowering plants pollinated by wild bees are in their majority weeds (93.1%). In our study system, *S. arvensis* and *D. carota* were the most abundantly pollinated floral resource. In line with our results, previous studies have already indicated that patches of wild vegetation in between the crops provide essential pollen resources when crops are not in flower [43] [44] [45] [46] [47] [37]. In fact, declines in pollinators are known to be linked to local extinctions of wild plants [40]. It is worth remarking that sunflower *H. annuus* turned out to be neither very abundant (5.7% reads) nor very common (9.1% FOO) floral resource within our study area. Sunflowers produce abundant pollen [48], which is easily available to certain groups of pollinators due the morphology of its flowers. However, this pollen is lower quality (low protein content) than that of many wild flowering plants [49], such as *P. rhoeas*, which is among the ten most pollinated plants by our wild bee communities [50]. Therefore, a diet relying heavily on sunflower pollen could have detrimental effects on wild bee fitness if it is not complemented with other pollen types [49]. Our ITS2 metabarcoding approach has successfully enabled us to precisely identify the highly diverse floral requirements of wild bees, supporting the idea that an abundant and diverse community of weeds make sunflower crops a higher quality habitat for wild bees. Hence, wild bees may be able to enhance agricultural production if the non-crop flowering plants around the agroecosystem are diverse enough to cover their feeding requirements [51].

### 4.3 Effects of flowering stage and farming system on the composition and diversity of the pollination spectrum of wild bees

Our metabarcoding data illustrates how environmental variables may have a strong influence on the selection of floral resource by wild bees. However, it also points out that the biodiversity metrics used, RRA and FOO, were not always perfectly congruent. For instance, the common sowthistle *Sonchus oleraceus* was the most visited plant during pre-flowering but not the most abundant species in terms of sequences. This is consistent with the fact that *S. oleraceus* produces low quantity of pollen during a short period (Percival, 1955; St John-Sweeting, 2011). Therefore, even if the flower is available in sunflower fields, wild bees may only collect small quantities of its pollen. This brings to light the need to consider not only the strong points of each metric but also their potential biases and assumptions (Deagle *et al*., 2019). While RRA provides a semi-quantitative measure of relative biomass (McClatchie and Dunford, 2003; Elbrecht and Leese, 2015), which could be a useful proxy of pollen biomass if treated with caution, FOO provides information about the low or high presence of the floral resource (Alberdi *et al*., 2018; Deagle *et al*., 2019). On one hand, RRA has shown a higher variance among laboratory and bioinformatic protocol used (Alberdi *et al*., 2018; Deagle *et al*., 2019) but it is more accurate and descriptive that FOO. On the other hand, FOO is more consistent, but the information it provides is less precise and this metric gives importance to rare taxa (Deagle *et al*., 2019). Combining RRA and FOO, after a robust laboratory and bioinformatic protocol (Alberdi *et al*., 2018), is potentially the best approach to accurately describe and monitor ecological communities (Deagle *et al*., 2019). In our study, both biodiversity metrics help in providing a global picture of the pollination spectrum of wild bees in sunflower crops.

Seasonal climatic variability is known to drive changes in pollination spectrum [52] [53]. Our results showed seasonal differences in the taxa of floral resources pollinated by wild bees when comparing pre-flowering and flowering seasons, which may be partly due to the phenology of flowering plants in our study area. In fact, we detected 11 plant species whose abundance in pollen loads varies significantly between both flowering periods. These plant species could be considered as indicators of a certain environmental condition, like for example, *D. carota* and *Daucus sp*. during sunflower flowering stage. Pollination niche breadth was significantly higher during sunflower pre-flowering season compared to the flowering season. Despite the imbalance in sample size (N_pre-flowering_ = 66, N_flowering_ = 119), a potential explanation could be the higher percentage of weeds flowering during the pre-flowering season. Indeed, more than half of the fields (54.6%) showed high density of weeds (>75% ground cover), while during flowering 63% of the fields had low density of weeds (<25% ground cover). Thus, we observed a congruence between the percentage of weeds covering the crop ground and the number of reads (RRA) detected in the pollen loads. Consistently, it has been reported that in more diverse environments, bees, especially social bees, not only increase the number of visitations of flowering plants [54] but also their intake of pollen (Kaluza *et al*., 2017, 2018). This goes in line with our results, which support the need of making weeds available for wild bees. Enhancing the biodiversity of weeds, would have a positive effect not only on wild bee fitness and social bee colony survival (Kaluza *et al*., 2017, 2018) but also on the productivity of pollinator-dependant crops, such as the sunflower [51]. Although the negative effects of agrochemicals on plant biodiversity have been widely reported [55] [56], our comparison of plant species diversity between organic and conventional crops, shows no statistical effect of the farming system. Even if the lack of agrochemicals is known to increase wild bee diversity [57], there may other factors influencing the number of plants available for the pollinators, such as for instance, landscape complexity, which has known to have a positive effect on pollinator-plant interactions [58]. Therefore, the landscape homogeneity occurring in intensive agricultural areas, such as LTSER ZA Plaine & Val de Sèvre, would be affecting the pollination spectrum in both, organic and conventional crops. In contrast, there is a compositional shift in terms of floral resources, as wild bees prefer different flowering plants depending on the farming system within the crop. For instance, *C. arvensis* is the most pollinated species in terms of abundance and an indicator plant species within conventional crops. Interestingly, this weed has been reported to be naturally tolerant to glyphosate [59], which is a widely used and non-selective herbicide [60]. Hence, weeds more pollinated within intensive crops may be resistant species to certain agrochemicals or the ones present only in organic fields could be more vulnerable to the use of agrochemicals. The differences in pollination spectrum could also be explained by the differences in wild bee communities inhabiting both types of crops. For instance, social bees were more often collected in conventional crops (79.8%) compared to organic crops (63.3%).

### 4.4 Body size and sociality as modulators of pollination spectrum in wild bees

Our findings highlighted a clear relation between wild bee body size on the pollination spectrum, which is consistent with previous studies. However, according to our ecological network wild bees of different size could pollinate similar taxa albeit with different intensity. This suggests that, even if wild bee body size had an effect, it would be in combination with other factors, such as social level as indicated through the significant interactions of PERMANOVA, CAP permutational test and GLM. Consistently, we found that in T1 most of wild bee species are social. In contrast, T3 shows the lowest diversity, almost the same number of bee species (6) and more than 75% of wild bee species are solitary (Table S1). This could be an effect of the low sample size of T3 (N = 15), compared to the other categories. However, this low diversity level makes sense as solitary bees tend to be more specialists [54] Kaluza *et al*., 2017, 2018). As shown in Figure S7, when comparing the most pollinated floral resources of the most abundant social wild bee *L. malachurum* with the most abundant solitary bee *A. flavipes*, we confirm the higher specialization of solitary bees compared to social bees as 67% of the pollination spectrum of *A. flavipes* was a single species, *S. arvensis*, while *L. malachurum* visited a higher number of plant species as shown by a more balanced percentage of read abundance. Moreover, this sheds light onto the fact that social bees are capable of foraging in different types of floral landscapes and thus may be more resistant to changes in flowering community composition. In contrast, certain solitary bees may depend on one or few specific plant species whose absence could lead to local extinctions of these bee species. There is, thus, an urge to manage agroecosystems in a way that they provide resources not only of social bees but also of solitary bees [54] Kaluza *et al*., 2017, 2018). This pollen metabarcoding study has enabled us to draw the broad picture of plant-wild bee interactions in an agroecosystem. Although our findings suggest some level of redundancy among the community of wild bees, several specific interactions illustrate the need for increased weeds diversity in agricultural landscapes to conserve a diverse community of wild bees.

## 5 Conclusion

Using an environmental (pollen) DNA approach, we characterized the pollination spectrum and shed light onto the interactions between wild bees, pollinated plants, environmental variables and wild bee species traits in sunflower crops. Despite a somewhat imbalance sampling, our study confirms that metabarcoding is an efficient tool to describe ecological interactions at the community level. Moreover, our study has brought to light hidden information about pollinated plants, which may not be available by conventional observational studies of visitation rates [15] [16]. In line with previous studies [39] [40] [41] [42], our work confirms the importance of weeds diversity for the conservation of wild bees in agricultural landscapes. The crop flowering stage and the farming system were shown to be strong drivers of change in the composition of plants pollinated by wild bees. The effect of crop flowering stage might be related to the seasonal climatic variability and the differences in the percentage of weeds available [52] [53]. In fact, the diversity of weeds available has been suggested to affect fitness and survival of wild bees (Kaluza *et al*., 2017, 2018). Although the composition of the community of pollinated plants vary according to the farming system, the diversity of plants pollinated is similar in organic and conventional fields. This may be due to the landscape homogenization within intensive agricultural areas [58], such as this case in the LTSER Zone Atelier Plaine & Val de Sèvre. Wild bee species traits, specifically body size and sociality, also impact the pollination spectrum, confirming results reported in previous studies [54] Kaluza *et al*., 2017, 2018). This work emphasizes the potential of eDNA to decipher plant-insect interactions and stresses the need to maintain weeds in intensive agroecosystems to conserve diverse communities of social and solitary wild bees in a context of global insect declines.

## 7 Data availability statement

The raw dataset generated and analyzed during the current study has been submitted to the NCBI Sequence Read Archive (SRA) under the BioProject PRJNA874523.

## 8 Author contributions

SB obtained the funding; SB and MQ designed the research; VB and SG provided the geographical and environmental data as well as logistic support for the fieldwork; MR identified the wild bee specimens; SB, PF, LM and MQ performed the fieldwork; PF and MQ carried out the laboratory work; MQ performed the statistical analysis; SB and MQ analyzed the data and wrote the manuscript; all authors contributed to revision of drafts and approved the final manuscript.

## 9 Acknowledgments

This work was supported by the ANR funded project IMAgHO (ANR-18-CE32-0002-01), the Government of Asturias, FICYT and “Plan de Ciencia, Tecnologia e Innovacion 2018-2022” from the Government of Asturias (Grant AYUD/2021/58607). We thank the contribution of Carolina Arteaga Garcia for designing and permitting us the use of the plant, wild bee and the logo original drawings included in the figures of this article and thank Irene Villalta for her help and advice in bee biology. Moreover, we thank Andreas Brachmann from the Sequencing Center of the Ludwig-Maximilian University (LMU) of Munich for his advice and efficiency in the library sequencing. Finally, we would like to thank the Mutualized Platform for Environmental Genomics of the Institut de Recherche sur la Biologie de l’Insecte (IRBI), UMR 7261 CNRS, for access to its facilities and logistic support.

