## Supplementary information for "Environmental variables and species traits as drivers of wild bee pollination in intensive agroecosystems -a metabarcoding approach"

**1. Extended protocol for metabarcoding laboratory analyses and bioinformatic**

*Wild bee sample processing, pollen DNA library preparation and flowering plant sequencing*

The washing protocol (Batuecas *et al.*, 2021) consisted in the immersion of each wild bee in 2 mL Tween®20 solution (0.1%) directly added to the 25 mL collection tube, which was then manually shaken during 2 minutes. Subsequently, wild bees were removed from the solution, rinsed with DNA-free water for 30 seconds and placed in a 1.5 mL Eppendorf tube for subsequent morphological species identification. Both, pollen solution and wild bee specimens were stored at -20ºC until further processing.

The DNA from pollen solutions was extracted using the epMotion 5075 workstation (Eppendorf, Hamburg, Germany) and the NucleoMag Tissue kit for purification from cells and tissue (Macherey-Nagel, Düren, Germany) following the manufacturer’s protocol. We included one extraction negative control and one PCR negative control were also included in the experiment. Initial pollen solution volume was 200 µl, the final elution was performed in 100 µl and the DNA extract was stored at -20 ºC until PCR amplification.

A fragment of 350 bp from the nuclear Internal Transcriber Spacer 2 (ITS2) DNA was amplified using the primer pair ITS2F (5′-ATGCGATACTTGGTGTGAAT-3′) and ITS4R (5′-TCCTCCGCTTATTGATATGC-3′) (White *et al.*, 1990; Chen *et al.*, 2010), correctly modified for high-throughput sequencing (HTS). PCR amplification reactions (25 µl) contained the following: 2 μl of template DNA, 1.5 μl of each primer [10 µM], 5 μl of 5X GoTaq (Promega) reaction buffer, 1 μl of MgCl2 [25 mM}, 1 μl of BSA [1 mg/ml], 0.5 μl of dNTPs [5 mM], 13.87 μl of molecular-grade water and 0.13 μl of GoTaq G2 Polymerase (Promega). PCR conditions were: 95 °C for 3 min, followed by 30 cycles of denaturation at 95 °C for 30 s, annealing at 55 °C for 30 s and elongation at 72 °C for 30 s, then a final elongation step was performed at 72 °C for 5 min (Batuecas *et al.*, 2021). Amplification success weas checked through a 2% gel electrophoresis. Successfully amplified amplicons were purified through a sodium acetate and ethanol precipitation protocol adding a 10:1 mix of ice cold 100% ethanol and 3M sodium acetate; 30 min centrifugation (4,000 rpm); adding ice cold 70% ethanol; 30 min centrifugation (4,000 rpm); drying at 80°C and resuspended in 40 μl molecular-grade water. Later, the ITS metabarcoding library was prepared by ligating Nextera XT indices through an eight cycle PCR (with the same conditions as for the initial PCR). The concentration of the successfully ligated samples (checked on agarose gel) was measured using a Qubit fluorometer (Life Technologies). Samples were then pooled equimolarly (100 ng each), selected by size (~534 bp) in a 1% agarose gel and purified using the GeneJet Gel Extraction kit (Life Technologies), according to manufacturer’s protocol and the pools eluted in 30 µl. Purified pools were combined into a 40 µl final pool [4 nM]. Sequencing runs were carried out on an Illumina Miseq using V2 chemistry (250 X 300 bp, 500 cycles) in the Sequencing Center within the Biozentrum of the Ludwig-Maximilian-University in Munich (Germany).

*ITS metabarcoding library filtering and taxonomic assignment*

The *FastQC* software (https://www.bioinformatics.babraham.ac.uk/projects/fastqc/) was used to check the quality of the libraries (demultiplexed *fastq* files) on forward and reverse reads. Pair of primers and 50 bp from reverse reads were removed using *cutadapt* (Martin, 2011) and merged with *PEAR* (Zhang *et al.*, 2014), setting Phred score 30 as a threshold. The subsequent quality filtering (*fastq_maxee* = 1), dereplication, denoising, insertion and deletions (indels) and chimera removal were performed using the *vsearch v2.8.2* software (Rognes *et al.*, 2016), which produced a *fasta* file containing Amplicon Single Variants (ASVs). These ASVs were clustered into Operational Taxonomic Units (OTUs) applying the centroid-based greedy clustering algorithm with a cut-off threshold of 97% (Xiong and Zhan, 2018) and a by-sample OTU table was mapped also using *vsearch* *v2.8.2* software (Rognes *et al.*, 2016). The obtained OTUs were classified against NCBI nr/nt using BLAST algorithm (Johnson *et al.*, 2008) and a customized script based on *rjson* (Couture-Beil and Couture-Beil, 2018) and *taxize* (Chamberlain and Szöcs, 2013) R packages in order to retrieve the best hit from the database. Among the taxonomic classification of the flowering plants, we discarded every OTU with a percentage of identity or a query coverage lower than 75% and 60%, respectively. OTUs with percentage of identity between 75% and 93% were assigned to family level, between 93% and 97% they were assigned to genus level and OTUs with more than 97% identity were initially assigned to species level. As an additional measure of accuracy, the taxonomic classification of each OTUs with a percentage of identity less than 100% was checked through a phylogenetic delimitation. To that purpose, all OTUs with less than 100% and with more than one species per genus, according to the list of weeds species present in the LTSER Zone Atelier Plaine & Val de Sèvre (SW France, Nouvelle-Aquitaine Region) (Gaba et al., 2010), were included in a phylogeny, together with all the species of the same genera whose sequences was available in NCBI nr/nt. The phylogenetic tree was constructed using a Maximum Likelihood approach (1,000 bootstraps) (Fig. S3) using the software IQ-TREE (Nguyen, Lam-Tung, et al., 2015). Moreover, reads of OTUs found in the negative controls were subtracted from all the samples and samples with less than 1,000 reads were removed for subsequent analyses.

**2. Detailed explanation of differences in pollination spectrum (plant taxa) among environmental variables and wild bee species traits**

At family level, there were differences in RRA between pre-flowering and flowering stages (Figure 2A, Table S2). For instance, while in pre-flowering stage, Brassicaceae was the most abundant family, during the flowering stage, Asteraceae comprised the highest number of reads. The most abundant species were *S. arvensis* during pre-flowering, and *D. carota* during flowering stage. In terms of occurrence (Table S2), Asteraceae was the most common family in both stages but the species with the highest FOO were *S. arvensis* (Brassicaceae) during pre-flowering and *D. carota* (Apiaceae) during flowering.

Regarding the type agriculture (Figure 2B, Table S2), Asteraceae was the family with the highest RRA in intensive agriculture fields, while Brassicaceae was the most abundant in biological crops. The family Convolvulaceae was 27 times abundant in intensive crops (RRA = 19.73%) than in biological crops (RRA = 0.77%). In fact, the plant species comprising the highest number of reads in intensive crops, *C. arvensis*, belongs to this family, while in crops with biological crops is *S. arvensis*. In terms of FOO (Table S2), Asteraceae was the most common family in both types of crops, while Convolvulaceae were the most represented in terms of DNA sequences, with higher rates in intensive crops compared to biological ones. The most common species in intensive crops was *S. arvensis* while in biological crops it was *D. carota*.

Pollen loads from small bees (T1 and T2) were characterized by high abundance of Asteraceae, while samples from medium-size bees (T3) displayed high abundance of Apiaceae, and larger bees (T4) had more reads from Brassicaceae (Figure 2C). At species level (Table S2), *D. carota* was the most abundant in T1 and T3 bees, while *P. hieracioides* and *S. arvensis* comprise the highest percentage of reads in T2 and T4 bees, respectively. In terms of FOO, Asteraceae was the most common family in all size categories but the most common species were *A. cotula* in T1, *D. carota* in T2 and T3 and *S. arvensis* in T4 bees. Alpha diversity also fluctuated among the four size categories (Figure 3C), with the highest value observed in T1, decreasing in T2, reaching its minimum in T3 and increasing again in T4. According to the Dunn test, the difference in alpha diversity between T1 and T3 bees was the only post-hoc comparison that was significant (K-W chi-squared = 2.459663, p-value = 0.0417).

With regards to sociality level, there were differences in the abundance of plant families detected from social and solitary bees (Figure 2 D). In social bees, the most abundant family was Asteraceae, while Brassicacea was the most represented in solitary bees. In addition, Convulvulaceae was much more represented in the pollination spectrum of social bees, while Apiaceae was more present in solitary bees. In both groups, the most represented plant species was *S. arvensis* (Table S2), followed in importance by *C. arvensis* in social bees and *D. carota* in solitary bees. FOO at family level showed Asteraceae as the most common family in both social levels, being, however, the most present species in both is *D. carota*, which belongs to Apiaceae.

**3. Supplementary figures**

**Figure S1.** A) Bar plot representing the number of collected specimens per wild bee species coloured by sociality. Species names are coloured by bee family and number of specimens of each category is written on the figure (N). B) Stacked bar plots of wild bee families classified according to flowering period. C) Stacked bar plots of wild bee families classified according to farming system.

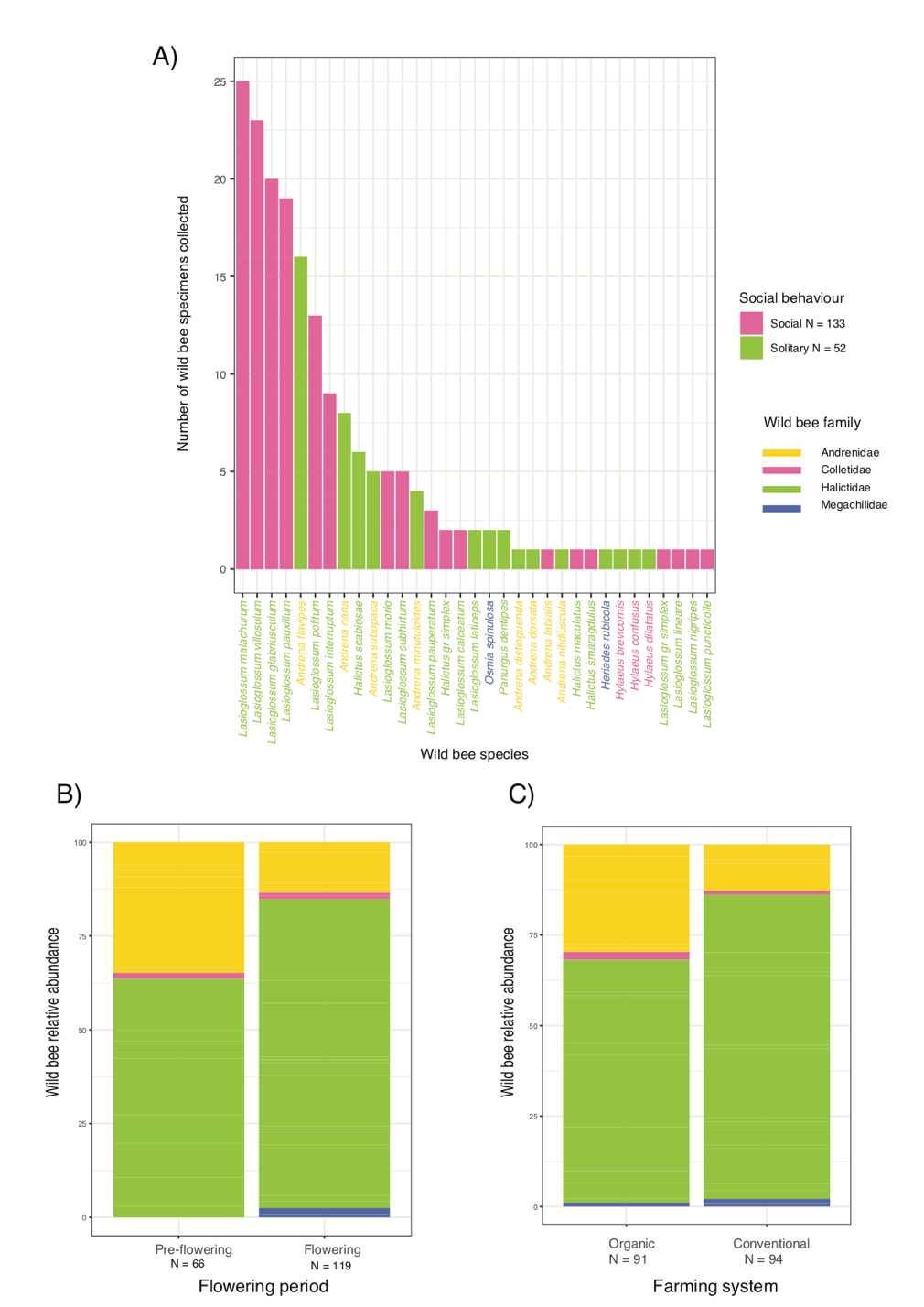

**Figure S2.** A) Accumulation curve as a proxy of completeness of the sampling representing the cumulative number of OTUs detected against the number of wild bee samples analyzed (n = 185). Horizontal solid line represents the number of OTUs expected with limitless sampling, based on bootstrapped estimates (999), B) Line plots representing rarefaction curves for each level of the studied environmental variables (flowering stage and farming system) and species traits (wild bee size and sociality).

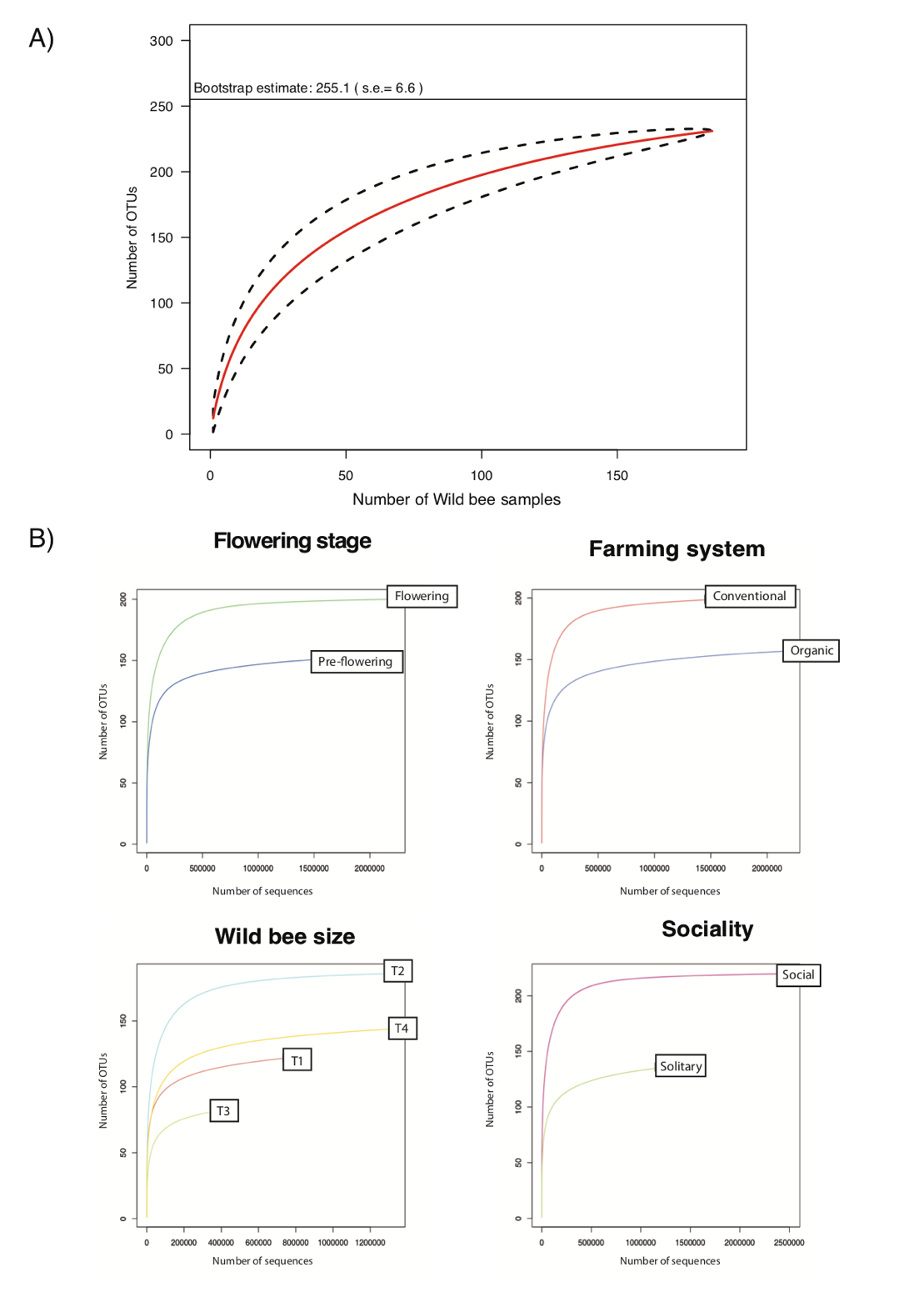

**Figure S3.** Phylogenetic tree (1,000 bootstraps) of the pollinated plant OTUs with percentage of identity higher than 97% but lower than 100%, and with more than one species within the same genus know to be present in the LTSER Zone Atelier Plaine & Val de Sèvre (SW France, Nouvelle-Aquitaine Region). This taxonomic phylogeny through midpoint rooting Maximum Likelihood (ML), uses reference sequences from GenBank (from which accession numbers are shown in the tips). Clades are coloured by plant family and bootstrap support are shown in the nodes.

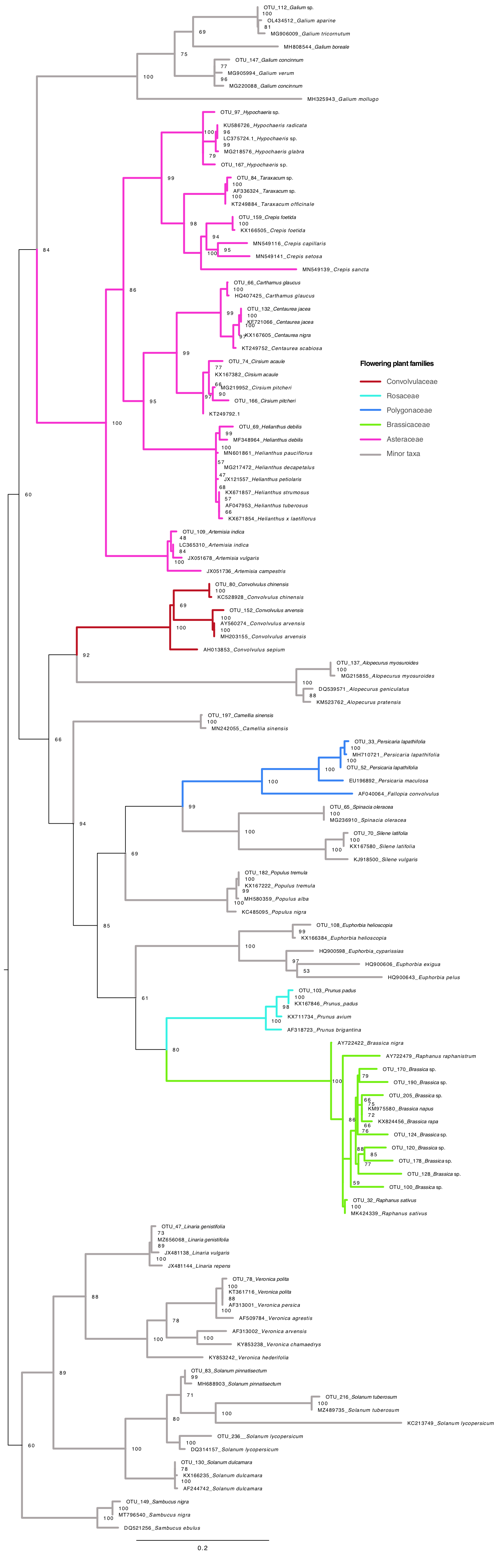

**Figure S4.** Boxplots representing the alpha diversity (exponential Shannon index) of flowering plants pollinated by wild bees in relation to A) sunflower flowering stage (pre-flowering vs flowering), B) farming system (organic vs intensive), C) Wild bee size categories (T1-T4) and D) sociality (social vs solitary bees). Significant variations are indicated within the plots (*) and Kruskal-Wallis test results are indicated on the top right of each plot.

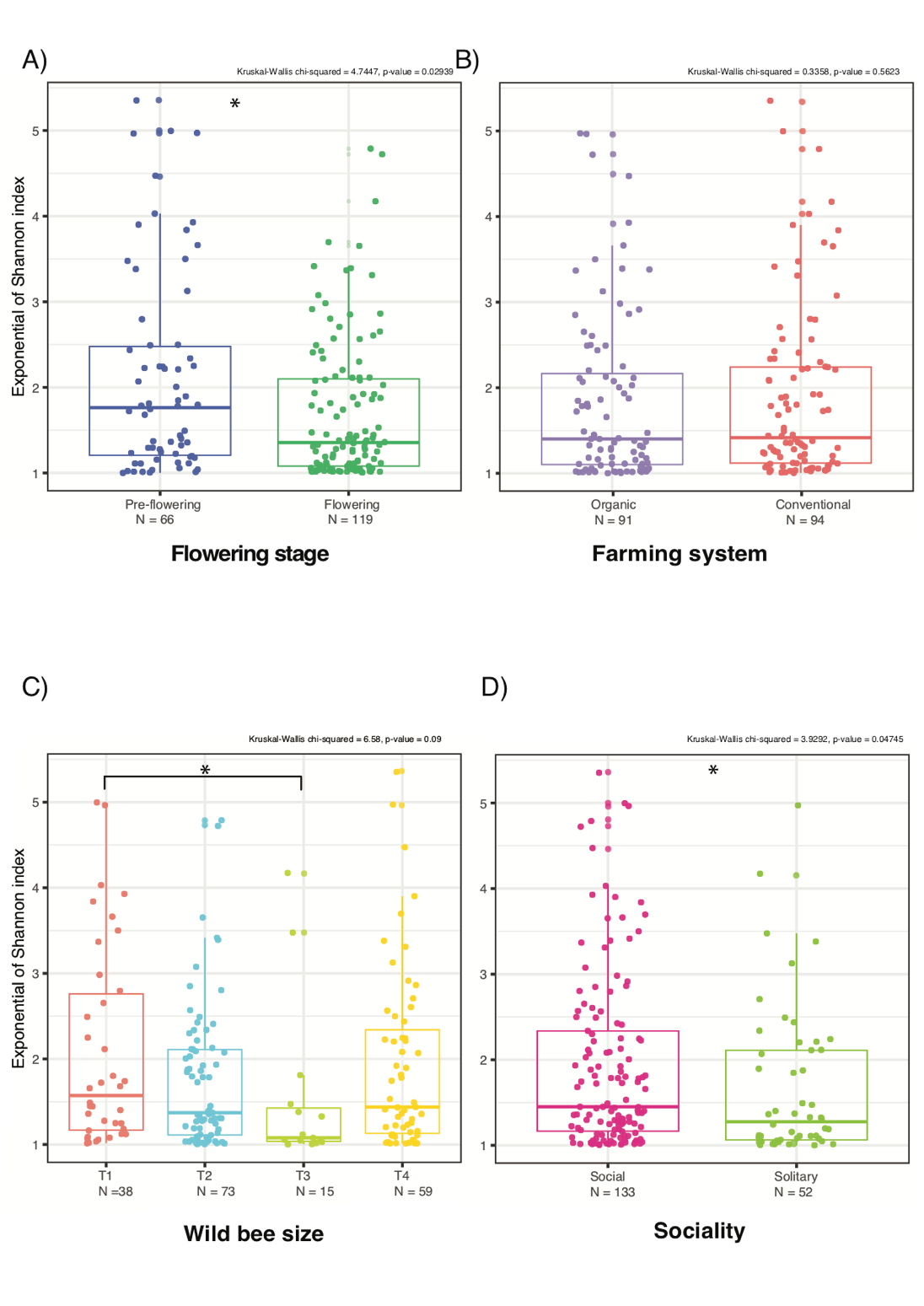

**Figure S5.** Principal Component Analysis (PCoA) based on Bray-Curtis distances at OTU level showing differences in community composition between A) sunflower flowering stage (pre-flowering vs flowering), B) farming system (organic vs intensive), C) Wild bee size categories (T1-T4) and D) sociality (social vs solitary bees). PERMANOVA test results are indicated on the top right of the plots.

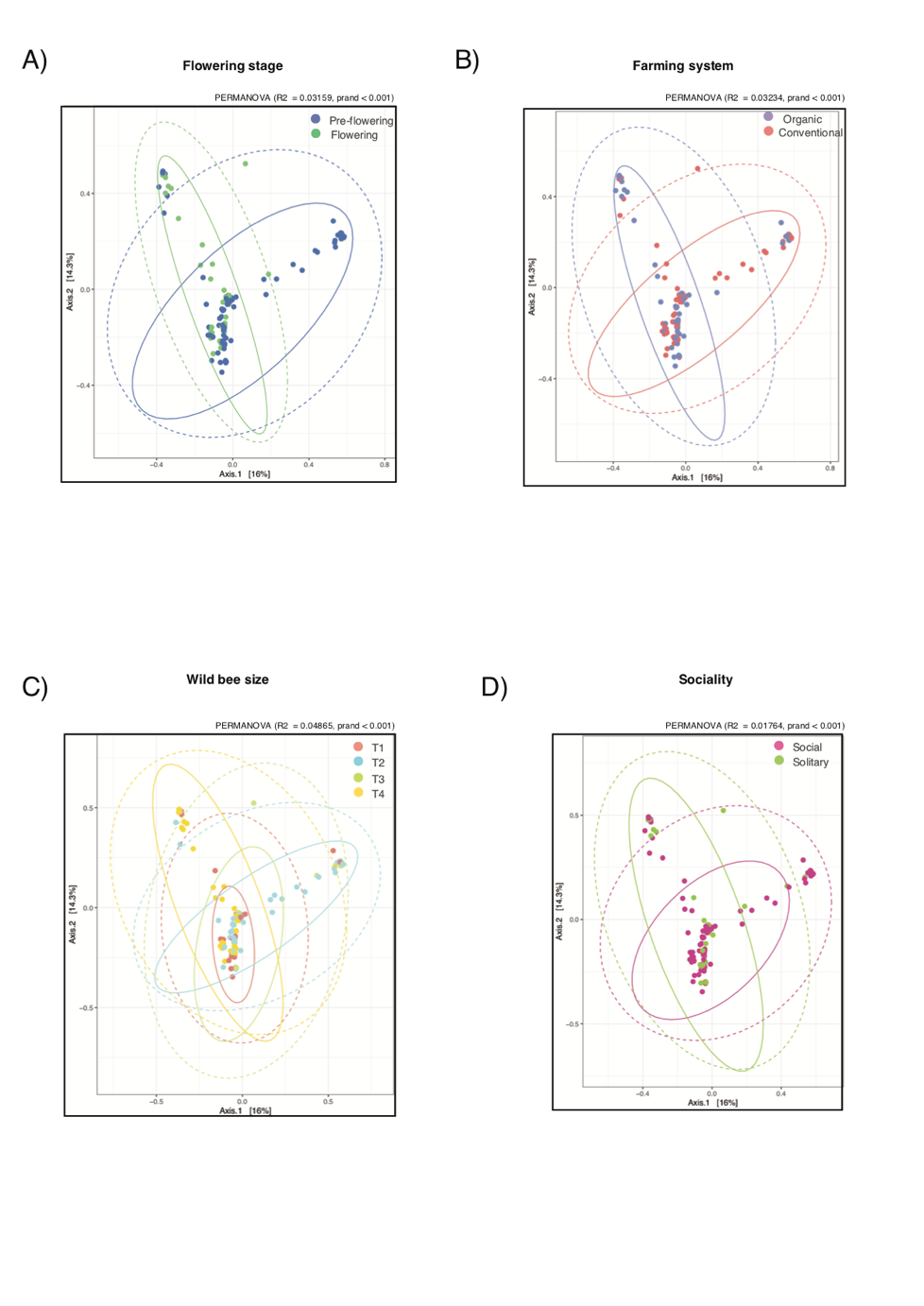

**Figure S6.** Community composition (beta diversity) visualized through Canonical Analysis of Principal coordinates (CAP) in relation to A) sunflower flowering stage (pre-flowering vs flowering), B) farming system (organic vs intensive), C) Wild bee size categories (T1-T4) and D) sociality (social vs solitary bees). CAP permutational test results are indicated on the top right of the plots.

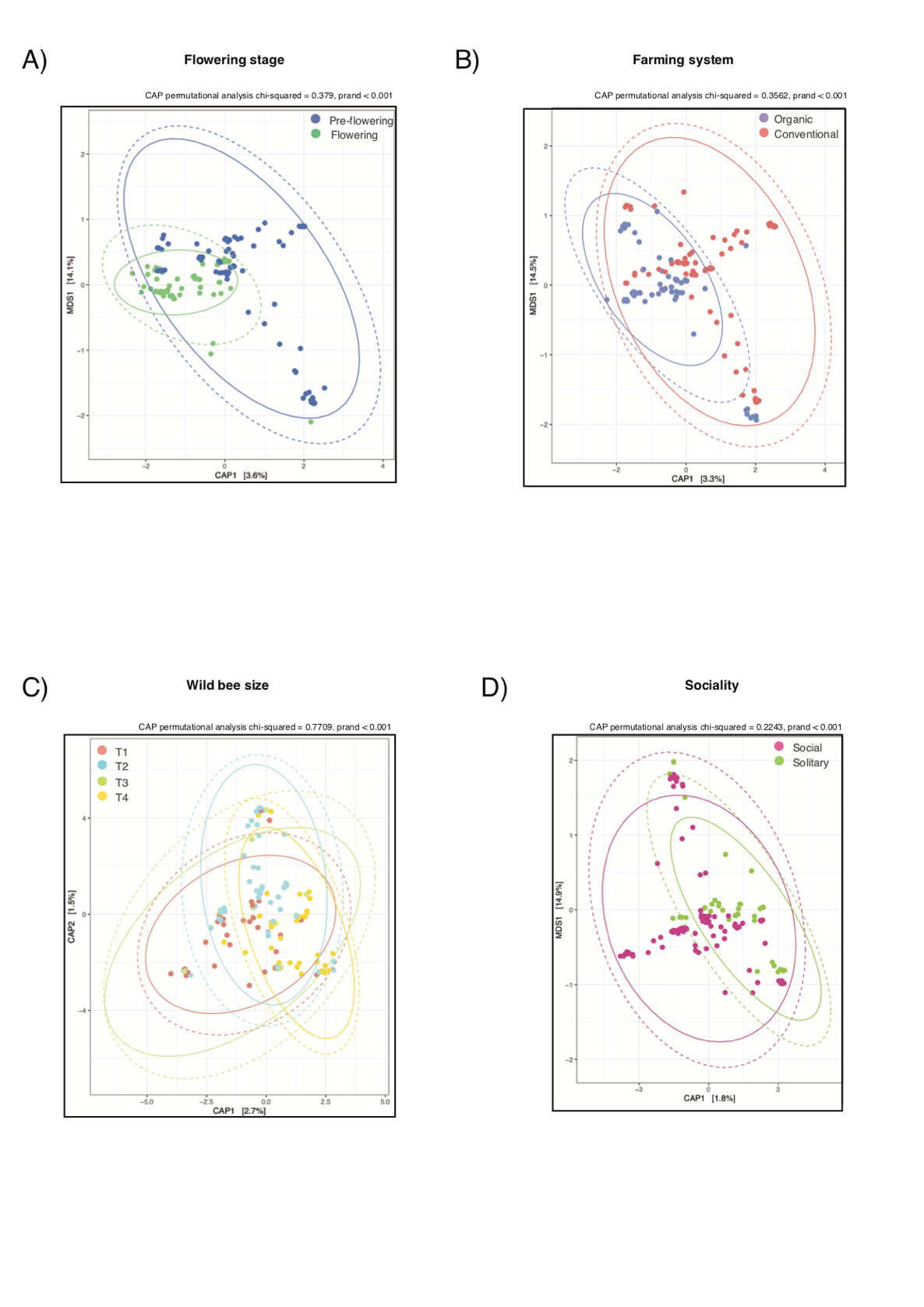

**Figure S7.** Relative Read Abundance (RRA) of the flowering plant species pollinated by the most abundant social species: the sharp-collared furrow bee *Lasioglossum malachurum*, and the most abundant solitary species: the yellow-legged mining bee *Andrena flavipes.*

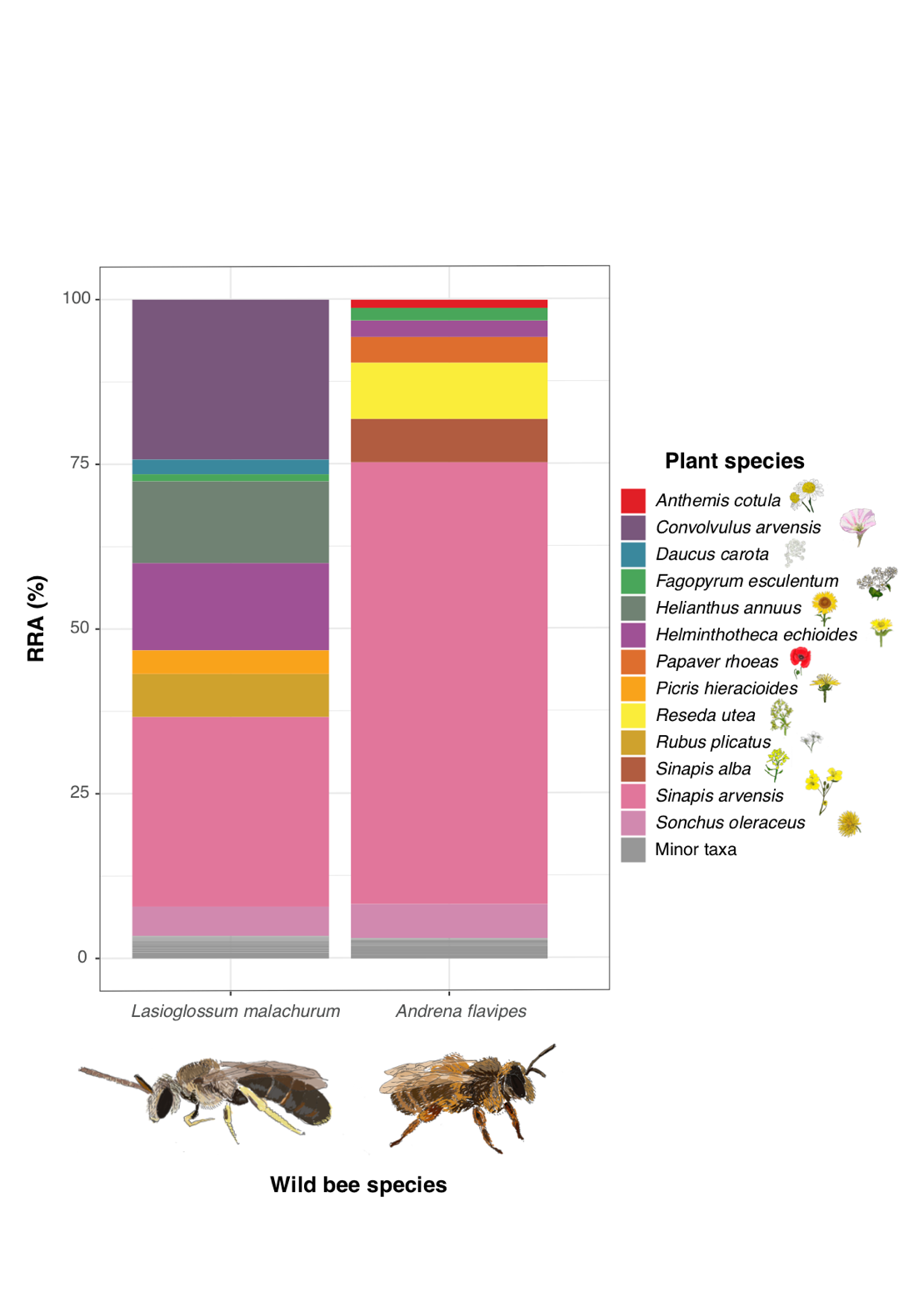

**Tables**

**Table S1.** A) List of wild bee samples used in this study, indicating the wild bee species, the sex, the wild bee body size, the code of the field where it was collected, the stage of the crop, the weather at the time of collection, the percentage of sunflower flowering, the percentage of weed cover, the sunflower height (cm) and the type of agriculture. B) Number of wild bee specimens per factor of environmental variable (flowering stage and farming system) and species trait (body size and sociality) selected to be included in the biodiversity analyses.

**Table S2.** Taxonomic assignment against NCBI of the 231 flowering plant Operational Taxonomic Units (OTUs) recovered from the present study, including the type of plant, the values of Relative Read Abundance (RRA) and the Frequency of Occurrence (FOO) for the complete sampling and for each level of the variables studied.

**Table S3.** List of OTUs that were differentially abundant in relation to at least one variable, their taxonomic classification, the Kruskal-Wallis Chi2 p-value and the environmental variable(s) or species trait(s) they are indicators of.

**Table S4.** Summary of the results of the statistical analyses, indicating the name of the analyses, the biodiversity metric tested, the variables included, the variable of the statistical indicator and the value of the p-value or prand.
